# Arteriolar degeneration and stiffness in cerebral amyloid angiopathy are linked to β-amyloid deposition and lysyl oxidase

**DOI:** 10.1101/2024.03.08.583563

**Authors:** Lissa Ventura-Antunes, Alex Nackenoff, Wilber Romero-Fernandez, Allison M Bosworth, Alex Prusky, Emmeline Wang, Cristian Carvajal-Tapia, Alena Shostak, Hannah Harmsen, Bret Mobley, Jose Maldonado, Elena Solopova, J. Caleb Snider, W. David Merryman, Ethan S Lippmann, Matthew Schrag

## Abstract

Cerebral amyloid angiopathy (CAA) is a vasculopathy characterized by vascular β-amyloid (Aβ) deposition on cerebral blood vessels. CAA is closely linked to Alzheimer’s disease (AD) and intracerebral hemorrhage. CAA is associated with the loss of autoregulation in the brain, vascular rupture, and cognitive decline. To assess morphological and molecular changes associated with the degeneration of penetrating arterioles in CAA, we analyzed post-mortem human brain tissue from 26 patients with mild, moderate, and severe CAA end neurological controls. The tissue was optically cleared for three-dimensional light sheet microscopy, and morphological features were quantified using surface volume rendering. We stained Aβ, vascular smooth muscle (VSM), lysyl oxidase (LOX), and vascular markers to visualize the relationship between degenerative morphological features, including vascular dilation, dolichoectasia (variability in lumenal diameter) and tortuosity, and the volumes of VSM, Aβ, and LOX in arterioles. Atomic force microscopy (AFM) was used to assess arteriolar wall stiffness, and we identified a pattern of morphological features associated with degenerating arterioles in the cortex. The volume of VSM associated with the arteriole was reduced by around 80% in arterioles with severe CAA and around 60% in cases with mild/moderate CAA. This loss of VSM correlated with increased arteriolar diameter and variability of diameter, suggesting VSM loss contributes to arteriolar laxity. These vascular morphological features correlated strongly with Aβ deposits. At sites of microhemorrhage, Aβ was consistently present, although the morphology of the deposits changed from the typical organized ring shape to sharply contoured shards with marked dilation of the vessel. AFM showed that arteriolar walls with CAA were more than 400% stiffer than those without CAA. Finally, we characterized the association of vascular degeneration with LOX, finding strong associations with VSM loss and vascular degeneration. These results show an association between vascular Aβ deposition, microvascular degeneration, and increased vascular stiffness, likely due to the combined effects of replacement of VSM by β-amyloid, cross-linking of extracellular matrices (ECM) by LOX, and possibly fibrosis. This advanced microscopic imaging study clarifies the association between Aβ deposition and vascular fragility. Restoration of physiologic ECM properties in penetrating arteries may yield a novel therapeutic strategy for CAA.

## Introduction

While neuritic plaques and neurofibrillary tangles are the hallmark lesions of Alzheimer’s disease (AD), vascular pathology has been recognized as an important contributor to AD for over 100 years^1,2^. In the 1930s, deposition of an amyloid material in the walls of cerebral and meningeal vessels was first reported, and two decades later, this phenomenon was linked to AD and termed congophilic angiopathy because the material was detected with a Congo red stain ^3,4^. Now known as cerebral amyloid angiopathy (CAA), this entity is defined by deposits of aggregated β-amyloid on cerebral blood vessels. Arterioles with CAA have disrupted architecture and a propensity toward hemorrhage, making it the second most common cause of intracerebral hemorrhage in the elderly after hypertension ^5-7^. CAA is more common with advancing age and frequently co-occurs with AD; more than 80% of patients with AD have some degree of CAA ^5,8^. CAA is linked to cognitive impairment independent of the association with AD ^9^. Several autosomal dominant variants of CAA are known, but most cases are sporadic. Greater clinical awareness of CAA began in the 1990s as magnetic resonance imaging (MRI) techniques were developed which could detect small areas of bleeding in the brain termed cerebral microhemorrhages (CMH). CMHs usually occur in a lobar distribution in CAA and are associated with white matter hyperintensities and enlarged perivascular spaces. This pattern has become the key diagnostic imaging feature of the disease ^10^. The recently updated Boston criteria provide excellent sensitivity and specificity for clinically diagnosing CAA ^11^.

Morphologically, β-amyloid forms deposits primarily in the tunica media layer of arterioles, and as vessels with CAA degenerate, they experience loss of VSM ^12,13^. This process mechanically impairs vascular reactivity, leading to suboptimal autoregulation of blood flow in the brain, ^14^ which has been seen in studies using arterial spin labeling (ASL) in patients with CAA as well as in mouse models ^13,15^. CAA is associated with the ε4 allele of apolipoprotein E (ApoE4) and has been reported to be associated with a coding variant in complement receptor 1 (CR1). Prior reports suggested late complement activation and various matrix metalloproteinases may play a role in vascular degeneration in CAA ^16,17^. Despite these molecular hints, surprisingly little is known about the mechanisms leading to vascular degeneration in CAA. Even the notion that β-amyloid plays a role in the progression of vascular degeneration has been questioned. Several authors independently reported that β-amyloid is not consistently present at sites of vascular rupture in CAA, creating doubts whether β-amyloid is a driving factor ^18,19^.

CAA is an important variable in experimental anti-β-amyloid immunotherapies for AD ^20^. These immunotherapies may be complicated by a side-effect currently termed ARIA or amyloid-related imaging abnormality ^21,22^. ARIA is marked by cerebral edema and/or microhemorrhages in a pattern that is typical of CAA, and the few pathological descriptions of ARIA invariably document the presence of CAA in affected patients ^23,24^. While most patients with ARIA have both imaging and symptomatic resolution, between 1 to 2% of patients in the clinical trials developed significant or life-threatening symptoms ^25^. This side effect may be due to rapid β-amyloid clearance from the brain through the perivascular pathway, leading to blood-brain barrier (BBB) dysfunction and inflammation. Because small vessel disease can independently lead to cognitive decline and may be an important variable in patient selection for emerging therapies, a better understanding of the mechanisms of vascular injury in CAA and identifying therapeutic targets that simultaneously address the parenchymal and vascular features of AD are important objectives.

To address gaps in the current understanding of how β-amyloid contributes to CAA pathology, we sought to characterize changes to vessel morphology associated with vascular degeneration and assess the degree to which β-amyloid deposition correlates with microvascular degeneration and microhemorrhage. We also interrogated the biophysical properties of vessels with β-amyloid deposits and assessed the degree to which the extracellular matrix crosslinking enzyme lysyl oxidase (LOX) is associated with vascular degeneration. Defining the cellular and molecular features of microvascular degeneration in CAA is important to understanding vascular dysfunction’s contribution to cognitive decline in AD and CAA.

## Materials and methods

Brain tissue for this analysis was obtained from clinical autopsies performed at Vanderbilt University Medical Center and the University of California Los Angeles Brain Bank (UCLA), including neurological controls and patients with CAA over a range of severities. For each subject, we selected tissue samples representing each cortical lobe (frontal, temporal, parietal, and occipital cortex), including any areas with visible microhemorrhages when they were present. When significant hemorrhages or stroke were present, the contralateral side was used for tissue acquisition. Cases were grouped according to the CAA severity reported in the autopsy, however, a much larger quantity of tissue was included in this study than was evaluated in the autopsy and because of the patchy nature of the disease, a wide range of severity was observable in most cases.

### Ethics statement

Brain tissue donation was approved by the institutional review board of Vanderbilt University Medical Center, and written informed consent was obtained from patients or their surrogate decision makers. All specimens used in the study were de-identified. The study received IRB oversight as part of our ongoing Observational Study of Cerebral Amyloid Angiopathy and Related disorders (OSCAAR, IRB# 180287). Deidentified tissue specimens sourced from the UCLA brain bank received similar institutional ethical oversight and were graciously provided by Dr. Harry V. Vinters.

### Tissue optical clearing

The CLARITY methodology consists of transformation of opaque biological tissues into a transparent hydrogel-tissue hybrid with preserved anatomical structure, proteins, and nucleic acids. The tissue becomes transparent after hydrogel infusion, which provides a support framework for the brain tissue, and lipid removal ^26^. We selected brain tissue samples of 1 to 3 cm thickness for tissue clearing (Fig. 1A). Fresh tissue blocks were incubated for 3-5 days in 4% paraformaldehyde (PFA), and embedded in an acrylamide hydrogel (4% acrylamide, 0,05% bis-acrylamide, 0.25 % temperature-triggering initiator VA-044 in 0.1 M phosphate-buffered saline in water-PBS). This study also used tissue samples that had been previously fixed and stored in 4% PFA. These samples were directly embedded with acrylamide hydrogel. Adequate tissue incorporation of the hydrogel requires an oxygen-free environment to prevent premature cross-linking of acrylamide. To accomplish this, we placed the tissue sample in a 10 mL centrifuge tube and added enough hydrogel to fill the tube, leveraging surface tension so the hydrogel protruded above the lip of the tube when it was sealed with wax with no trapped air bubbles. The tissue blocks remained in the hydrogel at 4°C for 3 weeks, then were moved to a 37°C water bath for 4 hours to activate the VA-044 acrylamide crosslinker and polymerize the hydrogel. After polymerization, the lipid component of the tissue was passively cleared with prolonged washing in a solution consisting of 0.2M boric acid and 4 % w/v sodium dodecyl sulfate (SDS) at pH 8.5 at 37°C with gentle agitation. The process of tissue clearing involved exchanging the buffer once or twice a week until the optimal translucence was achieved. The time required for the clearing process depended on the type of tissue being cleared. For fresh tissue, the process took between 8 to 12 weeks, but for previously fixed tissue and samples with severe CAA which has more blood in the parenchyma the process often took longer. Upon completion, we transferred the tissue into PBS with 0.1% Triton X-100 in a light booth equipped with a 1200W LED array (BESTVA DC Series) for 24-48 hours of exposure to intense visible spectrum light at 4°C to photo-depigment the tissue and suppress autofluorescence. The tissue was then washed and incubated in PBS with sodium azide 0.02% until staining.

**Figure 1:**
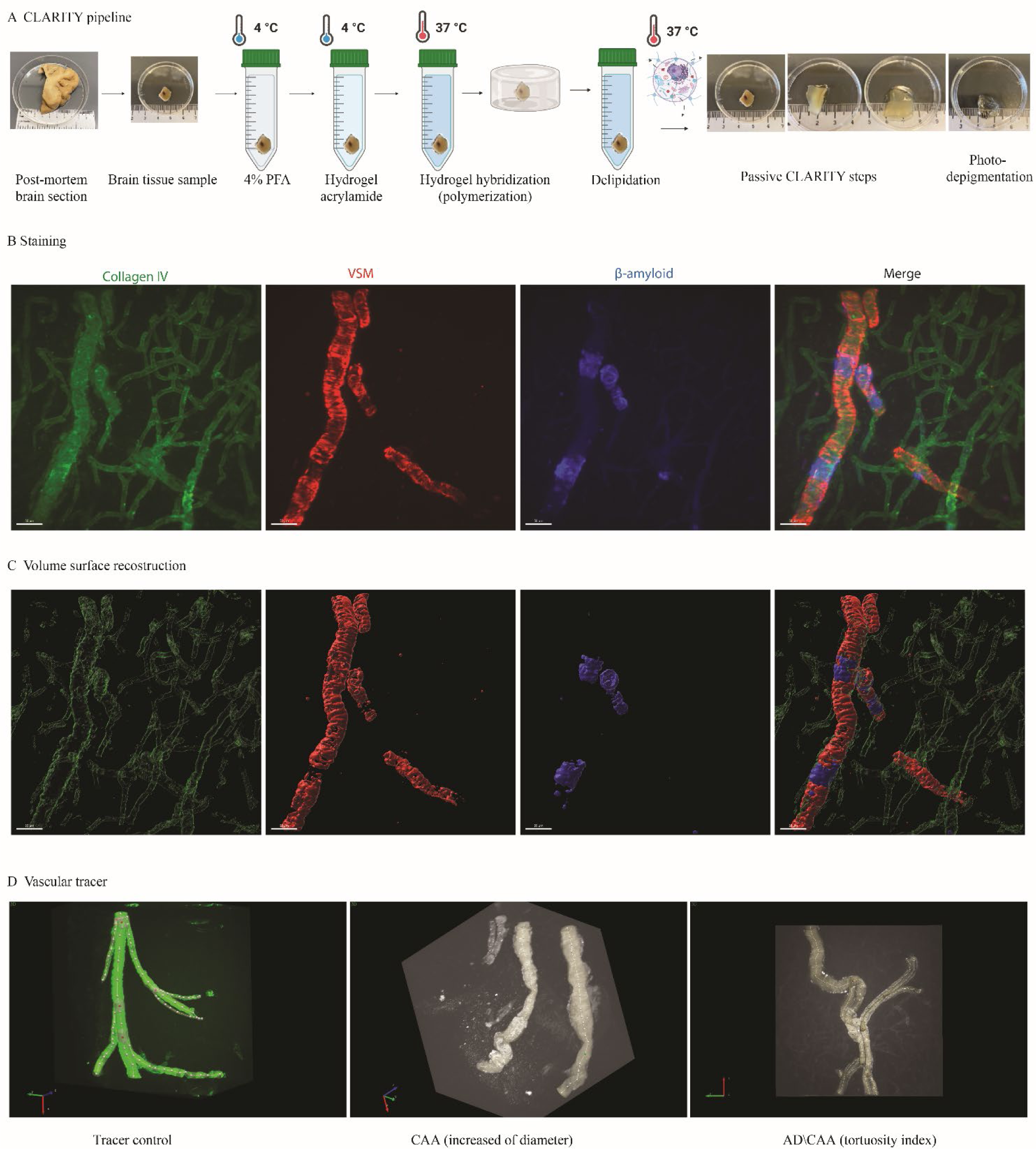
CLARITY methodology. **(A)** CLARITY methodology pipeline applied to human tissue samples, n=26 cases. **(B)** Immunostaining for vessels with Collagen IV-488 in green, vascular smooth muscle actin-Cy3 (VSM) in red, and methoxy-X04 to staining β-amyloid in blue. Scale bar with 100 µm. **(C)** Volume surface reconstruction from each of the images shown above using IMARIS software. Rotating three-dimensional versions of the images in B-C are also viewable in supplementary video 1. **(D)** We manually traced each penetrating arteriole using Vesselucida 360 on the smart manual mode. The first panel shows the tracing of a control vessel, the second and third panels show examples with CAA.

### Immunostaining

#### Staining for Lightsheet Microscopy

Optically cleared tissue blocks were washed overnight in PBS with 0.1% Triton X and preincubated with PBS containing 0.1% Triton X and 0.2M boric acid at 37°C. To stain the blood vessels, we utilized a directly conjugated primary antibody against collagen IV mouse anti-human, Alexa Fluor 488 (1:200, Cat# 53987182, Invitrogen) and incubated for 3 days in PBS with 0.1% Triton X. The tissue blocks were then washed and stained with conjugated antibodies for lysyl oxidase (5mg/ml, Cat# NB100-2527AF647 Lox antibody Alexa fluor 647, Novus Biologicals, LLC) and a mouse monoclonal antibody against VSM α-Smooth Muscle actin – Cy3 (5mg/ml, Cat# C6198, Sigma). Amyloid was stained with methoxy-X04 overnight (1:500 from in 5mM in distilled H2O, Cat # 4920, Tocris Bioscience) (fig. 1B, 5 and supplementary video 1), then washed in PBST 0.1% at 37°C.

#### Staining for Laser Scanning Confocal Microscopy

Floating sections of *post-mortem* human tissue were prepared on a Leica 1200 S vibratome with 50 μm and 100 μm thickness. β-amyloid was detected using thiazine red (1 µM, Cat# 2150-33-6, Chemsavers) associated with DAPI (0.5 µM, Cat# 1485, Vector lab) and a mouse monoclonal antibody against VSM α-smooth muscle actin (Cat# ABT1487, Millipore), associated with corresponding secondary antibody, Alexa Fluor 488 (Thermo Fisher A21206). Confocal images were acquired through the Vanderbilt Cell Imaging Shared Resource (CISR) using a Zeiss LSM 710 confocal laser-scanning microscope (Carl Zeiss, Germany) with a 40× objective.

### Lightsheet fluorescence microscopy (LSFM)

After immunostaining, optically cleared tissue samples were incubated in 68% thiodiethanol (TDE) (Cat# 166782, Sigma) overnight with a refractive index of 1.33 to match the optical diffraction of the microscope. Three-dimensional images were acquired with a Z1 lightsheet microscope (Carl Zeiss) at the Vanderbilt Cell Imaging Shared Resource (CISR) with 405 nm, 568 nm, 488 nm, and 638 nm lasers, using 20x objective illumination (Objective Clr Plan-Neofluor 20x/1.0 Corr nd=1.45 M32 85mm) and high-resolution scientific cameras that allow exceptional image quality and precise measurements (PCO. Edge sCMOS). Three-dimensional image stacks were assembled from plane-by-plane image collection in Z-plane depth between 750 – 2050 μm using ZEN software (Zeiss) for processing. In addition, larger scale images of vascular networks obtained from optically cleared tissue specimens (fig. 2A and video 2) were obtained on a SmartSPIM single-plane illumination light sheet microscope (LifeCanvas Technologies) with a 3.6× objective (NA = 0.2) and axially swept rolling shutter with 488 nm lasers. Following acquisition, images were transferred to a Dell Precision 7920 Tower with (2) Intel Xeon Gold 5218R CPU at 2.1 GHz and 526 GB of RAM running Windows 10 Pro for Workstations for stitching and processing. Images were stitched into Z-stacks with a modified version of Terastitcher Keyframe animations from the original image data were generated in Imaris (version 10, Bitplane, Oxford Instruments, USA).

**Figure 2:**
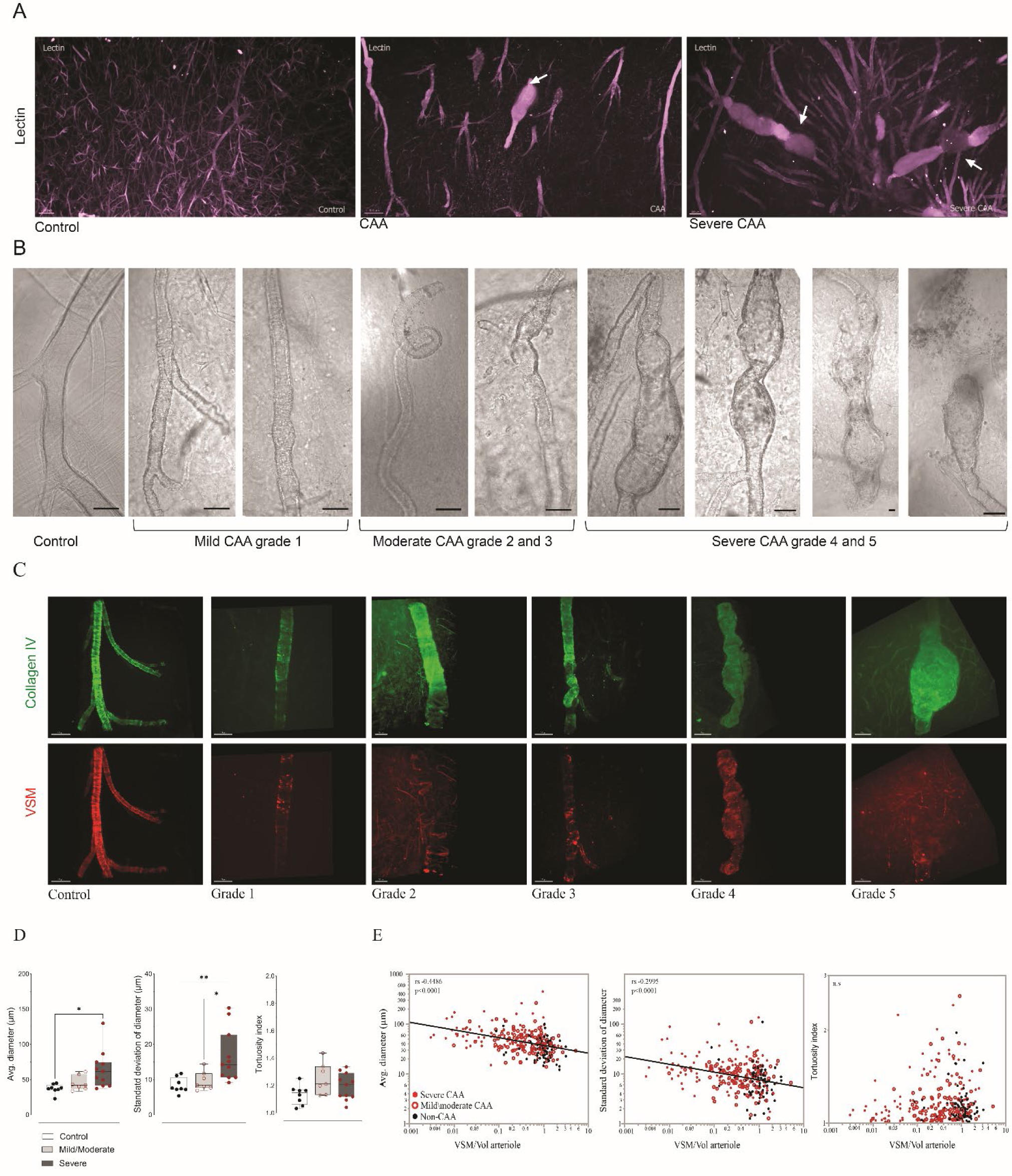
Morphological changes and grades of arterioles affected by CAA. **(A)** Macro view of vessels stained with tomato-lectin, in purple. Left panel, healthy cerebral vessels in a neurological control, with intact vascular wall morphology. Central and right panels, vessels from subjects with CAA show degenerative vascular morphology with enlarged vessel diameter, areas of distention and in the third image microhemorrhages (at arrows). Supplementary video 2 shows more details of these three views. **(B)** Phase-contrast imaging showing a range of levels of vascular degeneration in penetrating arterioles with CAA. **(C)** Fluorescent images showing the relationship between morphological degeneration (stained with collagen IV-488) and loss of VSM, in red. The control arteriole has a smooth vascular wall with intact VSM; mild, grade 1 degeneration is structurally intact with a thickened vessel wall; moderate degeneration, grades 2 or 3, has increased tortuosity and more pronounce VSM loss, and severe degeneration, grades 4 or 5 has progressively worsening dilation, areas of aneurysm or stricture and, at the terminal stage, vascular rupture. **(D)** Comparison of average diameter, variability of diameter (SD of diameter) and tortuosity of penetrating arterioles across cases (one-way ANOVA, *\* p>0.05.* **(E)** The Spearman correlation between average diameter, variability of diameter (SD), and tortuosity with VSM. Vascular dilation and the variability in the vascular diameter strongly correlate with VSM loss, tortuosity peaks at an early stage of vascular degeneration.

### Image analysis process and quantification

Images produced via LSFM were reconstructed and surface volume rendered using the software Imaris. We used automated detection of objects of interest to estimate features of vessel geometry, including surface volume, average diameter, and variability in diameter. Vascular β-amyloid volume, vascular LOX volume, and the volume of VSM were also estimated (fig. 1C). Image parameters used to measure the surface volumes included adjustments to account for immunostaining signal intensity including: (1) smoothing details set at 0.688, (2) thresholding adjusted to optimally recognize each voxel of immunostaining, and (3) rendering each structure volume (vascular volume, VSM volume, vascular β-amyloid, and vascular LOX) using the absolute intensity with background subtraction. Image size was approximately 30 GB with 16-bit images; size: X:1920 µm Y:1920 µm Z:800 µm to 900 µm. The Z size varied according to the object of interest, and the volume of the sample voxel size was around X 0.344 Y:0.344 Z: between 0.500 to 0.700. To trace vessels, we analyzed the three-dimensional images acquired at the LSFM using the Vesselucida 360 software (MBF Bioscience) to determine the vessels’ length, tortuosity, and diameter. The arterioles were traced using the smart manual mode (fig.1D). The Vesselucida 360 software was limited in its ability to trace severely damaged vessels, so those with rupture or significant discontinuity in staining were omitted from the assessment of tortuosity, and we estimated the length and volume of these arterioles with the IMARIS software using the detectable vascular fragments to estimate their geometry. For intact vessels, the software Vesselucida Explorer (MBF Bioscience) was used to calculate the number of segments, the average diameter, length, and tortuosity. The segment was modeled as a series of frusta, and the total length of the path is used to trace the segment. To estimate the average diameter, the software calculates the length-weighted mean. To estimate the tortuosity, the ratio of the actual length of the segment to the shortest distance between the endpoints of the segment is calculated so that a straight segment receives a value of 1, and as tortuosity increases and the segment assumes a more complex path to reach its destination, the value will increase, typically ranging as high as a value around 2.5.

### Atomic force microscopy (AFM)

AFM was performed on 30 µm thick post-mortem unfixed human tissue sections. Prior to imaging, sections were incubated with 1 µg/ml Hoechst (Cat# H1399, ThermoFisher) and 2 µM Thiazine red (Cat# 2150-33-6, Chemsavers) for 5 minutes at room temperature to label nuclei and amyloid aggregates, respectively (fig. 4). Sections were mounted on a Nikon Eclipse Ti microscope and submerged in PBS. Penetrating arterioles were identified through brightfield and fluorescence channels. The Bioscope Catalyst AFM was mounted onto the microscope platform to measure vessel stiffness. AFM probes (Bruker Nano MLCT-bio-drift compensated) with a spring constant of 0.03 N/m were calibrated and cross-sections through vessel walls were scanned at a frequency of 0.25 Hz in tapping mode, capturing scan areas of 4 to 10 µm within the vessel wall. A total of 6 vessels were measured from each case, selected from at least two different sections. Each scan consisted of 16,384 elastic modulus measurements which were averaged, and the resulting 6 averages were plotted as a box and whisker plot.

### Statistical analysis

The arteriolar surface volume, VSM, vascular β-amyloid and LOX volumes were calculated from surface rendering using IMARIS. Once the object surface was created, the volumes were recorded using automated tools as shown in fig. 1C. Vascular diameter, length, and tortuosity were generated by Vesselucida Explore. Because CAA is a patchy pathology, affecting some arterioles more than others even when they are in close proximity, for most analyses each penetrating arteriole was treated as an independent measurement. Correlations between vascular volume, LOX, β-amyloid and morphological features/tracing measurements were performed in JMP17 2.0 software using nonparametric Spearman correlation coefficients. P value <0.05 was considered statistically significant. In addition, one-way ANOVA was applied to the correlation between cases and average of diameter, SD of diameter and tortuosity. Comparison of the difference between CAA and non-CAA vascular stiffness was performed using Student t-test in GraphPad Prism software version 10 (GRAPH PAD software Inc, USA).

## Results

We analyzed 384 images from LSFM of 26 human post-mortem brains, including 11 cases with severe CAA, 7 cases with mild/moderate CAA (5 with concurrent AD, 2 with frontotemporal dementia), and 8 cases which served as neurological controls, without CAA, AD, or FTD. We performed three-dimensional microscopy on optically cleared tissue blocks and digitally created surface renderings of the images of cortical arterioles affected by different grades of CAA to evaluate morphological alterations in the vessels. This workflow is illustrated in fig. 1 and supplementary video 1.

Because of the uncertainty about whether β-amyloid is causative of vascular degeneration in CAA, we aimed to characterize the features of vascular degeneration in CAA agnostic to the presence of β-amyloid. Cortical arterioles from neurological control cases generally had smooth, thin, uniform vessel wall appearance, but we observed a range of morphological features in optically cleared specimens from patients with CAA which we felt were likely evidence of vascular degeneration (see the large-scale reconstruction of samples from a neurological control and two CAA cases shown in fig. 2A and in supplementary video 2). These features included vascular tortuosity, dilation of penetrating arterioles and dolichoectasia measured as the variability in the caliber of arterioles (with both strictures and aneurysms) and are illustrated in fig. 2A-B. We aimed to quantify these features and assess their association with VSM loss in the arteriolar wall, which is a well-established degenerative feature of CAA. Consistent with prior reports ^27^. Tortuosity, lumenal diameter, and variability in the lumen size were all detectable with LSFM using collagen IV staining as a primary marker (fig. 2C). These features seemed to represent a spectrum of vascular degeneration, which, for the purposes of our analyses, we defined in five grades ordered according to our perception of their severity. Healthy-appearing arterioles were designated as grade 0 (control). Arterioles with widening of the lumen but that remained straight with smooth contours were designated grade 1, then with mild dilation and increased tortuosity grade 2, then with increased variability in the vessel diameter and loss of the smooth vessel-wall contour and tortuosity grade 3, then with lumenal dilation and variability in lumen size grade 4, and finally, with the emergence of marked structural degeneration including aneurysm and rupture grade 5 (fig. 2B-C). We also found that CAA cases often have a wider arteriolar diameter compared with neurological control. The healthy arterioles had an average diameter of *36.40 ± 2.24 µm (mean ± sem)*, while the average diameter was *48.65 ± 3.33 µm* for mild/moderate CAA and *62.49 ± 4.32 µm* in donors with severe CAA (one-way ANOVA *F= 5.522, p=0.0110*). The variability in the diameter of penetrating arterioles was also greater in severe CAA compared with control and mild\moderate CAA (fig. 2D). To assess the validity of this grading scheme for vascular degeneration and ensure it correctly ordered the features of arteriolar degeneration, we evaluated the correlation between morphological features of degeneration with VSM loss. The loss of VSM from arterioles as CAA progresses is probably the most established and reliable marker of vascular degeneration in CAA at this time ^12,28^. Three-dimensional microscopy of control tissue showed arterioles had intact VSM covering a high percentage of the vessel wall, while the VSM signal was lost in vessels with CAA as the morphological alterations of the vascular wall progressed (fig. 2D and supplementary video 1), as others have previously reported ^29^. We hypothesized that the average arteriolar diameter would increase with the loss of VSM, and we found a strong inverse correlation between these two variables *(r_s_ (279) = ρ-0.4486, p<0.0001)*. Areas of stricture and dilation cause irregularity in the lumen which can be quantified as an increase in the variability in the average arteriolar diameter. We hypothesized that the variability in arteriolar diameter would increase with loss of VSM. Our analyses found that the variability in vessel diameter was inversely correlated with VSM volume *(r_s_ (279) = ρ-0.2995, p<0.0001)*. Finally, we hypothesized that tortuosity would peak at mild or moderate stages of VSM loss when pathology was focal and could exert asymmetric effects on the vessel wall. In both disease-free vessels and severely affected vessels with concentric pathology, we predicted the vessels would remain more-or-less straight. Our result showed that there was no significant correlation between tortuosity and VSM volume across the entire cohort, but when only cases with CAA were included, there was a weak correlation, peaking at an early disease stage *(r_s_ (236) = ρ 0.1787, p= 0.0096)*. Because tortuosity peaked at an early stage of VSM loss, we interpreted it as an early feature of vascular degeneration.

Next, we aimed to assess the degree to which vascular β-amyloid deposition correlated with morphological features of vascular degeneration. We stained vascular β-amyloid using Thiazine red in optically cleared tissue blocks and quantified the volume of amyloid (vascular Aβ volume) on penetrating arterioles in cortical specimens for comparison with the morphological features of degeneration identified in the first stage of this study. We also assessed staining of vascular β-amyloid deposition in tissue sections (fig. 3A) to compare with the overview of CLARITY three-dimension image (fig. 3E). Vascular β-amyloid volume strongly correlated with the loss of VSM across the cohort *(r_s_ (297) = ρ= −0.6612, p<0.0001)*, (fig. 3B). Spatial comparisons of the location of β-amyloid deposits and VSM staining consistently showed that VSM was lost from vessel segments with heavy β-amyloid deposition (see fig. 1B-C and supplementary video 1). In most affected vessels, β-amyloid formed a ring-shaped structure encircling the vessel wall within the tunica media and this ring in many cases had a central void which lacked Thiazine red staining. This central void occasionally stained for VSM (fig. 3C), but in most cases VSM staining was absent. In areas with patchy β-amyloid deposits, we observed that the rings of β-amyloid were in line with VSM staining. These observations led us to conclude the β-amyloid rings probably form around VSM cells which subsequently involute, a hypothesis supported by previous studies ^30^. We also observed occasional VSM staining in vessels with CAA outside of the tunica media which we hypothesized might be evidence of vascular fibrosis (see arrows in fig. 3C). In advanced CAA, the VSM layer is essentially replaced by β-amyloid.

**Figure 3:**
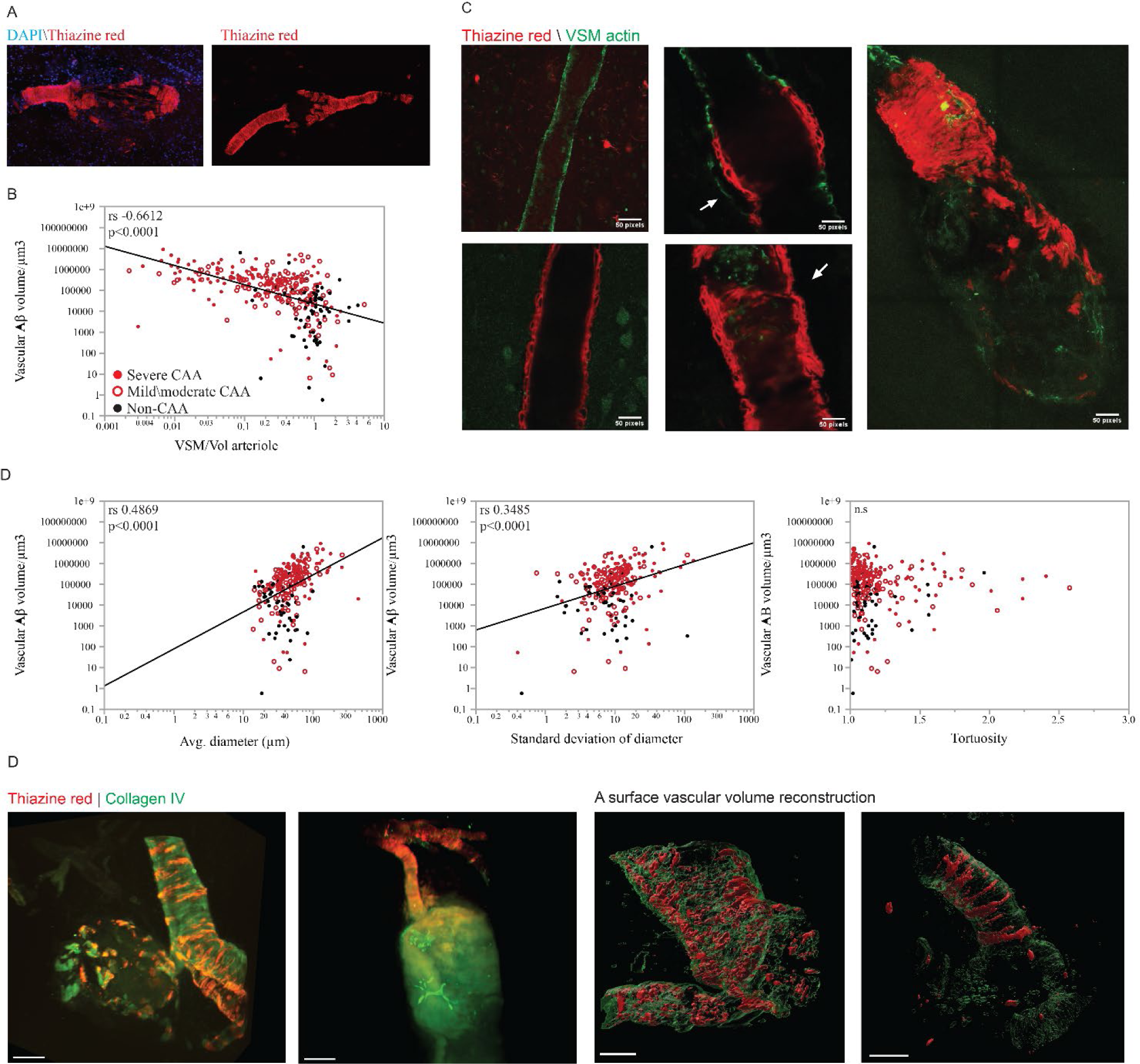
Association of β-amyloid accumulation in the wall of arteriole and structural degeneration and rupture. **(A)** Two-dimensional staining of β-amyloid with Thiazine red (red) and nuclei in DAPI (blue). **(B)** Spearman correlation between vascular β-amyloid surface reconstruction and VSM. **(C)** Two-dimensional staining of β-amyloid accumulation on arterioles and the absent staining of VSM in local β-amyloid accumulation. The arrow in the upper middle image shows VSM outside the tunica media consistent with a fibrotic process. The arrow in the lower middle image indicates the site of a focal stricture in the vessel wall. **(D)** Spearman correlation between vascular β-amyloid and average diameter, variability of diameter, and tortuosity. E. Three-dimensional images of ruptured arterioles with staining of β-amyloid with Thiazine red (red) and Collagen IV-488; the first two images show native fluorescence, the last two show vascular surface rendering. Disorganized “shards” of β-amyloid are present near the site of each rupture. Detailed views of these ruptured vessels can be seen in supplementary video 4.

Next, we assessed the relationship between vascular β-amyloid volume and vessel diameter, and we found they were strongly correlated *(r_s_ (241) = ρ= 0.4869, p<0.0001)*. Similarly, dolichoectasia, which we modeled as the variability in the diameter of vessels, correlated with vascular β-amyloid volume *(r_s_ (243) = ρ= 0.3485, p<0.0001)*, fig. 3D, suggesting that vessels lose their smooth barreled morphology as β-amyloid deposition progresses. Vascular β-amyloid volume did not correlate with tortuosity across the entire cohort but was modestly correlated with the index of tortuosity when vessels with CAA were evaluated alone *(r_s_ (196) = ρ= −0.1923, p=0.0038)*.

The tight correlation between vascular β-amyloid volume and degenerative morphological and cellular features in the arterioles strengthens the notion that β-amyloid likely plays a role in vascular degeneration. Several reports have called into question whether β-amyloid was present in vessels associated with microhemorrages, so we next attempted to address whether vascular β-amyloid is associated with microhemorrhage. We isolated 78 microhemorrhage images in our specimens which were visible on gross tissue inspection (either on natural or cut surfaces or after tissue clearance). These microhemorrhages ranged in size from a few hundred microns in diameter to several millimeters and appeared black, brown, or dark red. We also examined 13 severely dilated vessels which seemed near to rupture. In morphologically intact vessels with β-amyloid deposition, β-amyloid formed ring-shaped structures organized orthogonally to the direction of blood flow. In ruptured arterioles and those with severe focal dilations, the organized, ring-shaped deposits were disrupted into sharply contoured “shards” of β-amyloid. Others have described that β-amyloid deposits in CAA are rigid ^14^, so we interpreted this “shard-like” appearance as a shattering of the rings of β-amyloid. The expansion of the volume of these vascular segments could lead to the impression that β-amyloid deposition was proportionally decreased in traditional two-dimensional microscopy, but in every microhemorrhage we imaged β-amyloid was present. Fig. 3E and supplementary video 4 show ruptured arterioles and shards of β-amyloid deposits in CAA vessels in three-dimensional images co-stained with Thiazine red (in red) and Collagen IV (vascular arteriole in green).

The “shard-like” pattern of β-amyloid deposits in vessels at late stages of degeneration and in microhemorrhages suggests that vessels in CAA may become brittle which could contribute to their propensity toward rupture. To assess whether cortical arterioles with β-amyloid deposition are stiffer than healthy vessels, we performed AFM on post-mortem brain tissue from a subset of our cases: 10 cases (5 cases with CAA and 5 controls, table 1). Hoechst and brightfield imaging associated with thiazine red were used to identify vessels which had been cut cross-sectionally during tissue sectioning and stiffness measurements were made across the cut-surface of the vessel wall. Using AFM, we found that vessels containing heavy β-amyloid deposits were significantly stiffer than vessels in control tissue, with mean stiffness values of CAA vessels ranging from 167 to 175 kPa while control vessels ranged from 36 to 68 kPa. Vessels with CAA were on the order of four-fold stiffer than vessels from donors without CAA (Fig. 4A). The average AFM measurement per case in non-CAA *(40.81±7.19)* and CAA *(157.4 ±6.25)* is shown in Fig. 4B. We can infer from these results that structural changes in the arteriolar wall are occurring in tandem with β-amyloid accumulation which almost certainly impact the vessel’s function, both in terms of its ability to autoregulate and maintain smooth blood flow.

**Figure 4:**
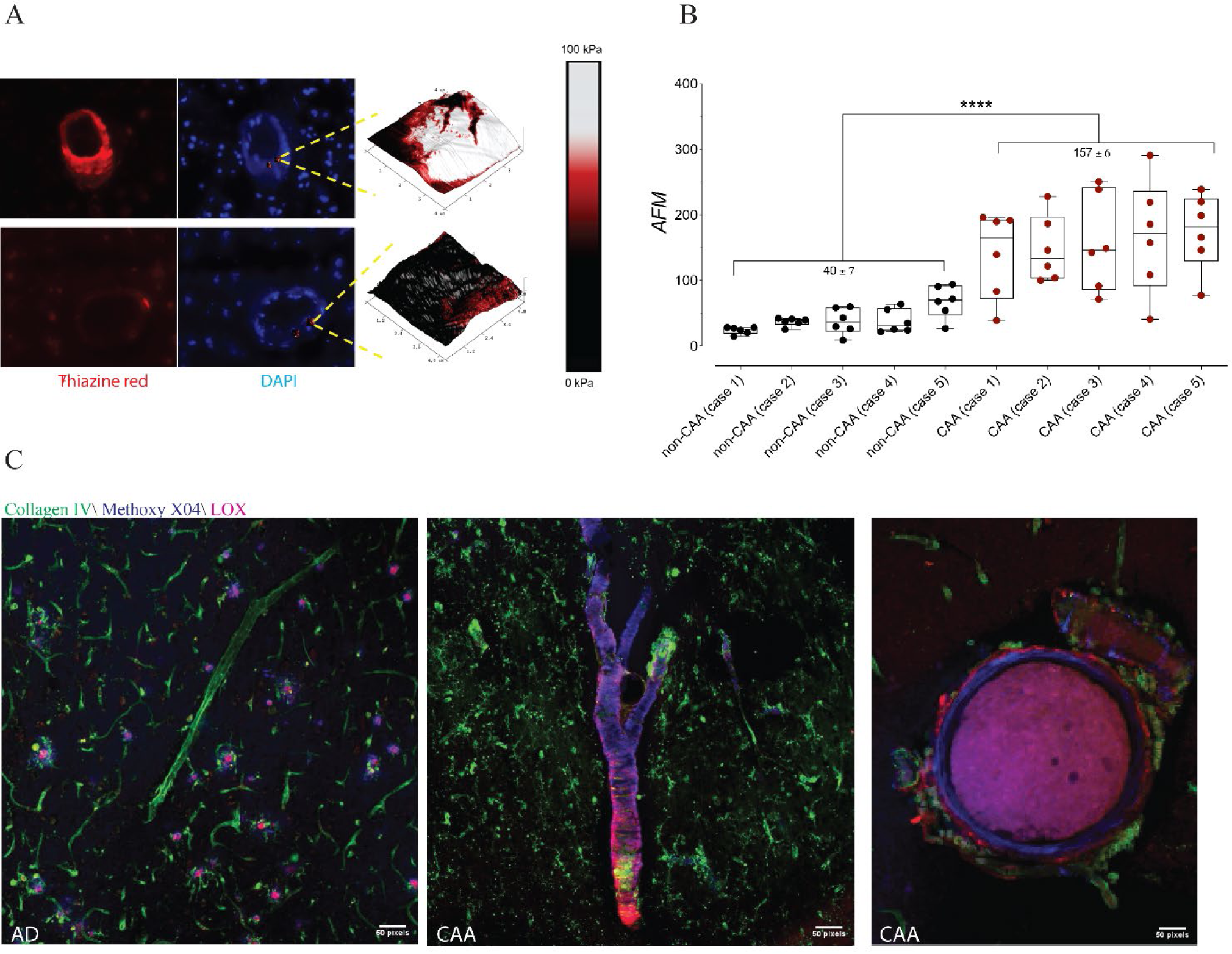
Atomic force microscopy (AFM) and the stiffness of arterioles. **(A)** Coronally sectioned profiles of arterioles from cases with CAA and without CAA were stained with thiazine red and DAPI. Cross-sections of the vessel wall were evaluated by AFM to determine tissue stiffness (example density maps across the vessel wall are shown for each condition). **(B)** Average stiffness in kPa for non-CAA (40.81±7.19) and CAA cases (157.4 ±6.25) are shown. **(C)** Increased tissue stiffness could be partly mediated by the activity of extracellular matrix crosslinking enzymes like lysyl oxidase (LOX). Confocal microscopy images of arterioles stained with collagen IV-488 (green), LOX (magenta), and β-amyloid staining with methoxy X04 (blue) shows substantial LOX deposition in CAA, especially in degenerating arterioles (rightmost image).

**Table 1.** Cases information and pathology description.

| **Category** | **Case ID** | **Age/**<br>**Sex** | **Neuropathology** | **Cause of death** | **PMI** |
| --- | --- | --- | --- | --- | --- |
| Severe CAA | CAA 1 | 90+M | Severe CAA, focal cerebellar CAA. A1B1C0. Rare microinfarcts. | Bronchopneumonia / sepsis | 48h |
| Severe CAA | CAA 2 / AFM case 6 | 62M | Intracerebral hemorrhage and severe CAA. APOE 3/4, A0B1C0. | Right frontal and left parietal hemorrhage | 35h |
| Severe CAA | CAA 3 | 86M | Intracerebral hemorrhage, severe CAA, severe AD – A3B3C3. | Intracerebral hemorrhage and pulmonary embolus | 18h |
| Severe CAA | CAA 4 / AFM case 8 | 71M | Intracerebral hemorrhage, severe CAA, severe AD – A3B3C3, remote right parietal infarct. | Right frontal hemorrhage | 12h |
| Severe CAA | CAA 5 / AFM case 9 | 81M | 3cm right frontal hemorrhage, severe CAA, AD pathology A2B1C2. | Urinary tract infection, in hospice care for end stage dementia | 19h |
| Severe CAA | CAA 6 / AFM case 7 | 68F | Severe CAA including cerebellum, severe AD – A3B3C3. | Hospice care for end-stage dementia | 9h |
| Severe CAA | CAA 7 | 61M | Severe CAA, AD pathology A2B3C3, numerous microinfarctions, subarachnoid siderosis | Intracerebral hemorrhage | 10h |
| Severe CAA | CAA 8 | 69M | Severe CAA, no Alzheimer's pathology. Scattered infarcts. | Sepsis, heart disease | 18h |
| Severe CAA | CAA 9 | 75F | Severe CAA, AD pathology A3B1C1. | GI infection (in hospice care). Remote episode of inflammatory CAA | 12h |
| Severe CAA | CAA 10 | 75M | Severe CAA, AD pathology A3B1C1. Subarachnoid siderosis, microinfarctions | Intracerebral hemorrhage | 38h |
| Severe CAA | CAA 11 | 81F | Severe CAA, Braak stage VI plaque pathology | End stage dementia | 21h |
| Severe CAA | AFM case 10 | 65F | Severe CAA, Braak stage VI plaque pathology | End stage dementia | 19h |
| Mild/mod CAA | CAA/AD 1 | 81F | Moderate CAA, severe AD pathology A1B3C3. Scattered cerebral infarctions | Heart disease and severe dementia | 36h |
| Mild/mod CAA | CAA/AD 2 | 68F | Mild/moderate CAA, severe AD – A3B3C3 | In hospice care for end-stage dementia | 15h |
| Mild/mod CAA | CAA/AD 3 | 67F | Moderate (focally severe) CAA, Braak stage VI plaque pathology | End-stage dementia | 28h |
| Mild/mod CAA | CAA/AD 4 | 66M | Moderate CAA, severe AD – A3B3C3. Brainstem predominant Lewy body disease | End-stage dementia in hospice care | 50h |
| Mild/mod CAA | CAA/mixed dementia 5 | 82M | Mild/moderate CAA. AD pathology – A3B0C2. LATE (TDP-43 stage 3/3). Diffuse Lewy body disease, hippocampal sclerosis | End-stage dementia in hospice care. | 8h |
| Mild/mod CAA | CAA/FTD 6 | 66M | Moderate CAA. Frontotemporal lobar degeneration due to TDP43 (AD pathology scoring A3B0C0). | Behavioral variant frontotemporal dementia | 10h |
| Mild/mod CAA | CAA/FTD 7 | 71M | Mild/moderate CAA. Frontotemporal lobar degeneration due to TDP43 (AD pathology scoring A3B1C0). | Corticobasal degeneration in hospice care | 6h |
| No CAA | Control 1 | 62M | Craniotomy site from remote subdural hemorrhage, microinfarction. | Portal vein thrombosis, sepsis | 11h |
| No CAA | Control 2 | 45M | Intracerebral atherosclerosis | Basilar artery stroke | 13h |
| No CAA | Control 3 | 87M | Left frontoparietal lacunar infarct, scattered rare neuritic plaques. | Myocardial infarction | 12h |
| No CAA | Control 4 | 87M | Left MCA stroke, Alzheimer's pathology scoring A1B1C0. | Stroke (contralateral hemisphere used in study) | 6h |
| No CAA | Control 5 / AFM case 5 | 66F | Vertebrobasilar atherosclerosis with pontine and cerebellar infarcts. | Pulmonary emboli, stroke. | 33h |
| No CAA | Control 6 | 72M | Left cerebellar lacunar infarction | Acute respiratory distress syndrome, heart failure. | 12h |
| No CAA | Control 7 | 68M | None (Alzheimer's pathology scoring A1B0C0) | Cardiogenic shock from intraoperative bleeding. | 24h |
| No CAA | Control 8 | 66M | None | Pneumonia and sepsis | 24h |
| No CAA | AFM case 1 | 65F | Lewy Body Disease, no CAA | End-stage dementia | 17h |
| No CAA | AFM case 2 | 27M | None | Complications of pulmonary hypertension | 11h |
| No CAA | AFM case 3 | 78M | Braak stage III/IV, no CAA | unknown | 20h |
| No CAA | AFM case 4 | 90+F | Braak stage VI, no CAA | End stage dementia | 16h |

Total of 26 cases included 11 CAA cases, 5 AD with mild CAA, 2 FTD with mild CAA, and 8 non-CAA cases without AD, or FTD. Five additional cases are included (4 non-CAA and 1 with CAA) which were included in the group assessed with atomic force microscopy, but were not included primary analysis for the other figures due to the volume of available tissue.\* PMI: post-mortem interval. Cerebral amyloid angiopathy (CAA); Alzheimer's disease (AD); Atomic force microscopy (AFM).

**Table 2.** Average of vascular tracing.

| **Cases** | **CAA grades** | **Tortuosity** | **SD of Diameter** | **Avg. Diameter $\mu\text{m}$** | **Vessel length $\mu\text{m}$** |
| --- | --- | --- | --- | --- | --- |
| Severe CAA 1 | Grade 3 | 1.228 | 11.269 | 41.245 | 1184.5428 |
| Severe CAA 2 | Grade 2 and 3 | 1.292 | 15.098 | 86.509 | 1446.7313 |
| Severe CAA 3 | Grade 4 and 5 | 1.313 | 14.255 | 74.757 | 1353.1763 |
| Severe CAA 4 | Grade 2 and 3 | 1.126 | 10.621 | 52.572 | 1189.6472 |
| Severe CAA 5 | Grade 5 | 1.095 | 22.708 | 65.047 | 1094.6128 |
| Severe CAA 6 | Grade 3 | 1.207 | 9.150 | 37.361 | 721.23543 |
| Severe CAA 7 | Grade 2 and 3 | 1.128 | 10.615 | 41.025 | 1936.4815 |
| Severe CAA 8 | Grade 2 and 3 | 1.337 | 30.335 | 72.428 | 1117.2443 |
| Severe CAA 9 | Grade 3 | 1.242 | 16.422 | 49.626 | 1550.9087 |
| Severe CAA 10 | Grade 1 and 2 | 1.119 | 13.538 | 61.611 | 1270.6133 |
| Severe CAA 11 | Grade 1 and 2 | 1.041 | 28.627 | 129.898 | 1373.4366 |
| CAA/AD 1 | Grade 2 | 1.129 | 8.399 | 42.125 | 1311.4108 |
| CAA/AD 2 | Grade 1 | 1.212 | 7.687 | 37.266 | 1468.3261 |
| CAA/AD 3 | Grade 3 | 1.340 | 10.374 | 43.079 | 1808.241 |
| CAA/AD 4 | Grade 1 | 1.437 | 11.766 | 61.280 | 1166.5144 |
| CAA/mixed 5 | Grade 3 | 1.199 | 7.998 | 39.992 | 1038.9729 |
| CAA/FTD 6 | Grade 1 and 3 | 1.133 | 14.421 | 57.440 | 1543.2614 |
| CAA/FTD 7 | Grade 1 and 3 | 1.305 | 6.921 | 33.566 | 1947.288 |
| Control 1 | Grade 0 | 1.252 | 7.388 | 44.835 | 1564.0481 |
| Control 2 | Grade 0 | 1.132 | 7.200 | 22.931 | 1033.110 |
| Control 3 | Grade 0 | 1.165 | 11.127 | 35.236 | 1508.5254 |
| Control 4 | Grade 0 | 1.052 | 5.399 | 43.644 | 944.1893 |
| Control 5 | Grade 0 | 1.171 | 11.725 | 38.542 | 1490.8495 |
| Control 6 | Grade 0 | 1.033 | 7.174 | 35.993 | 985.287 |
| Control 7 | Grade 0 | 1.163 | 7.597 | 36.852 | 1512.4164 |
| Control 8 | Grade 0 | 1.092 | 9.128 | 38.453 | 1613.9934 |

Average per case of morphological level (described in figure 2), average of diameter and length.

**Table 3.** Average vascular volumes of β-amyloid, lysyl oxidase, and vascular smooth muscle per patient.

| *Category* | *Case#* | *% Aβ/Arteriole* | *% Lox/Arteriole* | *% VSM/Arteriole* |
| --- | --- | --- | --- | --- |
| *Severe CAA1* | CAA1 | 15.08 ± 3.10% | 13.20 ± 4.54% | 39.7 ± 8.07% |
| *Severe CAA 2* | CAA 2 | 42.60 ± 5.54% | 27.70 ± 5.73% | 11.02 ± 2.89% |
| *Severe CAA 3* | CAA 3 | 13.89 ± 3.34% | 17.91 ± 4.58% | 47.59 ± 5.46% |
| *Severe CAA 4* | CAA 4 | 10.46 ± 3.68% | 6.00 ± 3.44% | 47.41 ± 8.42% |
| *Severe CAA 5* | CAA 5 | 26.94 ± 4.07% | 8.26 ± 2.15% | 14.52 ± 3.78% |
| *Severe CAA 6* | CAA 6 | 1.48 ± 1.10% | 6.74 ± 5.65% | 86.62 ± 2.85% |
| *Severe CAA 7* | CAA 7 | 65.11 ± 9.71% | 0.93 ± 0.52% | 16.95 ± 7.83% |
| *Severe CAA 8* | CAA 8 | 17.05 ± 6.66% | 3.17 ± 3.06% | 44.75 ± 13.66% |
| *Severe CAA 9* | CAA 9 | 13.91 ± 5.96% | 2.52 ± 1.23% | 49.72 ± 10.42% |
| *Severe CAA 10* | CAA 10 | 64.37 ± 4.77% | 23.27 ± 6.11% | 10.52 ± 4.72% |
| *Severe CAA 11* | CAA 11 | 23.72 ± 8.70% | 2.50 ± 1.30% | 21.21 ± 8.25% |
| *CAA/AD 1* | AD 1 | 16.97 ± 3.02 % | 13.17 ± 3.20% | 37.91 ± 5.70% |
| *CAA/AD 2* | AD 2 | 0.99 ± 0.65% | 0.23 ± 0.23% | 73.40 ± 6.76% |
| *CAA/AD 3* | AD 3 | 23.56±8.69% | 0.52±0.51% | 0.20 ± 0.14% |
| *CAA/AD 4* | AD 4 | 19.01 ± 7.12% | 14.34 ± 6.26% | 25.03 ± 5.56% |
| *CAA/mixed 5* | AD 5 | 25.74 ± 3.62% | 8.63 ± 1.99% | 22.00 ± 2.96% |
| *CAA/FTD 6* | FTD 1 | 23.73 ± 7.91% | 9.84 ± 5.37% | 38.58 ± 8.23% |
| *CAA/FTD 7* | FTD 2 | 21.79 ± 9.97% | 0.63 ± 0.37% | 38.44 ± 9.21% |
| *Control* | Control 1 | 3.91 ± 12.80% | 0.01 ± 0.01% | 73.67 ± 9.29% |
| *Control* | Control 2 | 0.01 ± 0.00% | 0.00 ± 0.00% | 78.47 ± 8.86% |
| *Control* | Control 3 | 13.50 ± 2.73% | 0.94 ± 0.31% | 57.53 ± 4.85% |
| *Control* | Control 4 | 1.68 ± 0.56% | 0.21 ± 07% | 79.71 ± 5.86% |
| *Control* | Control 5 | 10.01 ± 3.49% | 4.08 ± 3.34% | 44.61 ± 5.91% |
| *Control* | Control 6 | 13.14 ± 3.20% | 14.03 ± 7.27% | 4.84 ± 0.43% |
| *Control* | Control 7 | 0.13 ± 0.08% | 0.49 ± 0.15% | 64.91 ± 5.75% |
| *Control* | Control 8 | 1.06 ± 0.57% | 0.08 ± 0.02% | 102.38 ± 22.74% |

Data is presented as percent coverage (volume/vascular volume) on penetrating arterioles. The values correspond to the average per case.

**Table 4.**
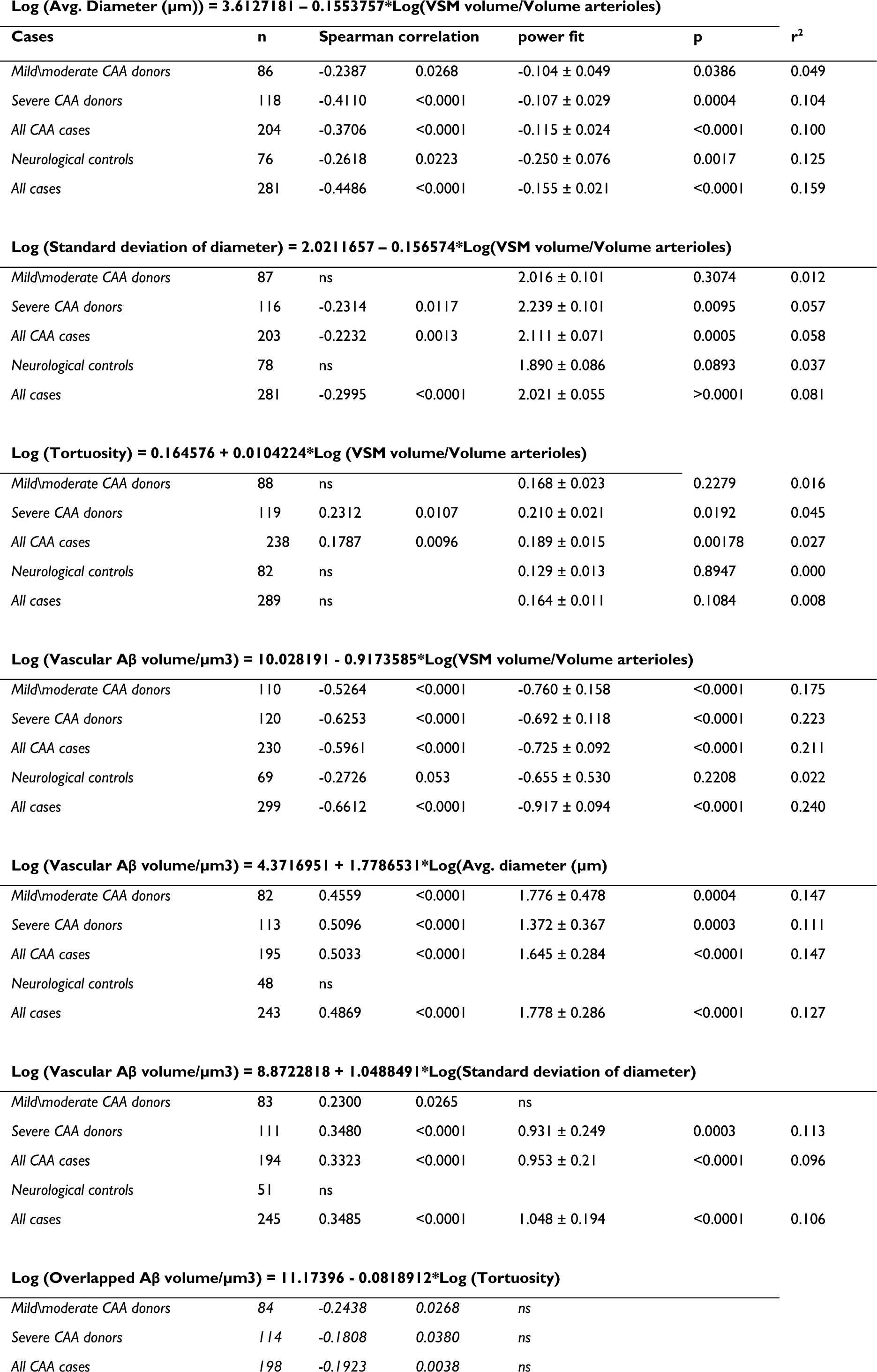
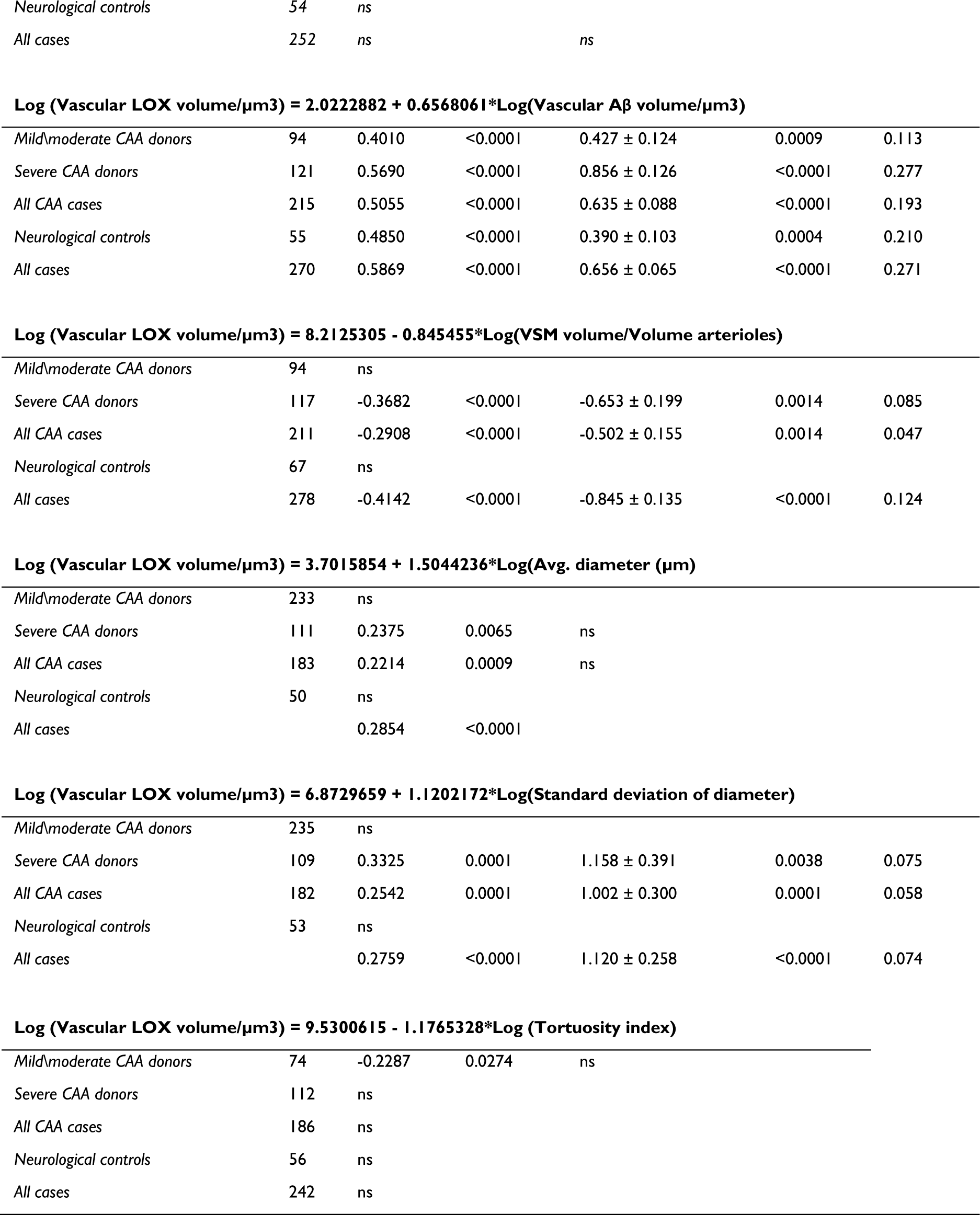
Spearman correlation detail for all observation Log (Avg. Diameter (µm)) = 3.6127181 – 0.1553757*Log(VSM volume/Volume arterioles)

| Cases | n | Spearman correlation |  | power fit | p | r<sup>2</sup> |
| --- | --- | --- | --- | --- | --- | --- |
| Mild/moderate CAA donors | 86 | -0.2387 | 0.0268 | -0.104 ± 0.049 | 0.0386 | 0.049 |
| Severe CAA donors | 118 | -0.4110 | <0.0001 | -0.107 ± 0.029 | 0.0004 | 0.104 |
| All CAA cases | 204 | -0.3706 | <0.0001 | -0.115 ± 0.024 | <0.0001 | 0.100 |
| Neurological controls | 76 | -0.2618 | 0.0223 | -0.250 ± 0.076 | 0.0017 | 0.125 |
| All cases | 281 | -0.4486 | <0.0001 | -0.155 ± 0.021 | <0.0001 | 0.159 |
| $$\text{Log (Standard deviation of diameter)} = 2.0211657 - 0.156574 * \text{Log(VSM volume/Volume arterioles)}$$ |  |  |  |  |  |  |
| Mild/moderate CAA donors | 87 | ns |  | 2.016 ± 0.101 | 0.3074 | 0.012 |
| Severe CAA donors | 116 | -0.2314 | 0.0117 | 2.239 ± 0.101 | 0.0095 | 0.057 |
| All CAA cases | 203 | -0.2232 | 0.0013 | 2.111 ± 0.071 | 0.0005 | 0.058 |
| Neurological controls | 78 | ns |  | 1.890 ± 0.086 | 0.0893 | 0.037 |
| All cases | 281 | -0.2995 | <0.0001 | 2.021 ± 0.055 | >0.0001 | 0.081 |
| $$\text{Log (Tortuosity)} = 0.164576 + 0.0104224 * \text{Log (VSM volume/Volume arterioles)}$$ |  |  |  |  |  |  |
| Mild/moderate CAA donors | 88 | ns |  | 0.168 ± 0.023 | 0.2279 | 0.016 |
| Severe CAA donors | 119 | 0.2312 | 0.0107 | 0.210 ± 0.021 | 0.0192 | 0.045 |
| All CAA cases | 238 | 0.1787 | 0.0096 | 0.189 ± 0.015 | 0.00178 | 0.027 |
| Neurological controls | 82 | ns |  | 0.129 ± 0.013 | 0.8947 | 0.000 |
| All cases | 289 | ns |  | 0.164 ± 0.011 | 0.1084 | 0.008 |
| Mild/moderate CAA donors | 110 | -0.5264 | <0.0001 | -0.760 ± 0.158 | <0.0001 | 0.175 |
| Severe CAA donors | 120 | -0.6253 | <0.0001 | -0.692 ± 0.118 | <0.0001 | 0.223 |
| All CAA cases | 230 | -0.5961 | <0.0001 | -0.725 ± 0.092 | <0.0001 | 0.211 |
| Neurological controls | 69 | -0.2726 | 0.053 | -0.655 ± 0.530 | 0.2208 | 0.022 |
| All cases | 299 | -0.6612 | <0.0001 | -0.917 ± 0.094 | <0.0001 | 0.240 |
| Mild/moderate CAA donors | 82 | 0.4559 | <0.0001 | 1.776 ± 0.478 | 0.0004 | 0.147 |
| Severe CAA donors | 113 | 0.5096 | <0.0001 | 1.372 ± 0.367 | 0.0003 | 0.111 |
| All CAA cases | 195 | 0.5033 | <0.0001 | 1.645 ± 0.284 | <0.0001 | 0.147 |
| Neurological controls | 48 | ns |  |  |  |  |
| All cases | 243 | 0.4869 | <0.0001 | 1.778 ± 0.286 | <0.0001 | 0.127 |
| Mild/moderate CAA donors | 83 | 0.2300 | 0.0265 | ns |  |  |
| Severe CAA donors | 111 | 0.3480 | <0.0001 | 0.931 ± 0.249 | 0.0003 | 0.113 |
| All CAA cases | 194 | 0.3323 | <0.0001 | 0.953 ± 0.21 | <0.0001 | 0.096 |
| Neurological controls | 51 | ns |  |  |  |  |
| All cases | 245 | 0.3485 | <0.0001 | 1.048 ± 0.194 | <0.0001 | 0.106 |

$$\text{Log (Overlapped A}\beta\text{ volume}/\mu\text{m}^3) = 11.17396 - 0.0818912 * \text{Log (Tortuosity)}$$
|  |  |  |  |  |
| --- | --- | --- | --- | --- |
| Mild/moderate CAA donors | 84 | -0.2438 | 0.0268 | ns |
| Severe CAA donors | 114 | -0.1808 | 0.0380 | ns |
| All CAA cases | 198 | -0.1923 | 0.0038 | ns |

|  |  |  |  |  |
| --- | --- | --- | --- | --- |
| Neurological controls | 54 | ns |  |  |
| All cases | 252 | ns |  | ns |
| Mild/moderate CAA donors | 94 | 0.4010 | <0.0001 | 0.427 ± 0.124 | 0.0009 | 0.113 |
| Severe CAA donors | 121 | 0.5690 | <0.0001 | 0.856 ± 0.126 | <0.0001 | 0.277 |
| All CAA cases | 215 | 0.5055 | <0.0001 | 0.635 ± 0.088 | <0.0001 | 0.193 |
| Neurological controls | 55 | 0.4850 | <0.0001 | 0.390 ± 0.103 | 0.0004 | 0.210 |
| All cases | 270 | 0.5869 | <0.0001 | 0.656 ± 0.065 | <0.0001 | 0.271 |
| $$\text{Log (Vascular LOX volume/}\mu\text{m}^3) = 8.2125305 - 0.845455 * \text{Log(VSM volume/Volume arterioles)}$$ |  |  |  |  |  |  |
| Mild/moderate CAA donors | 94 | ns |  |  |  |  |
| Severe CAA donors | 117 | -0.3682 | <0.0001 | -0.653 ± 0.199 | 0.0014 | 0.085 |
| All CAA cases | 211 | -0.2908 | <0.0001 | -0.502 ± 0.155 | 0.0014 | 0.047 |
| Neurological controls | 67 | ns |  |  |  |  |
| All cases | 278 | -0.4142 | <0.0001 | -0.845 ± 0.135 | <0.0001 | 0.124 |

$$\text{Log (Vascular LOX volume/}\mu\text{m}^3) = 3.7015854 + 1.5044236 * \text{Log(Avg. diameter (}\mu\text{m))}$$
|  |  |  |  |  |
| --- | --- | --- | --- | --- |
| Mild/moderate CAA donors | 233 | ns |  |  |
| Severe CAA donors | 111 | 0.2375 | 0.0065 | ns |
| All CAA cases | 183 | 0.2214 | 0.0009 | ns |
| Neurological controls | 50 | ns |  |  |
| All cases |  | 0.2854 | <0.0001 |  |
| $$\text{Log (Vascular LOX volume/}\mu\text{m}^3) = 6.8729659 + 1.1202172 * \text{Log(Standard deviation of diameter)}$$ |  |  |  |  |  |  |
| Mild/moderate CAA donors | 235 | ns |  |  |  |  |
| Severe CAA donors | 109 | 0.3325 | 0.0001 | 1.158 ± 0.391 | 0.0038 | 0.075 |
| All CAA cases | 182 | 0.2542 | 0.0001 | 1.002 ± 0.300 | 0.0001 | 0.058 |
| Neurological controls | 53 | ns |  |  |  |  |
| All cases |  | 0.2759 | <0.0001 | 1.120 ± 0.258 | <0.0001 | 0.074 |

$$\text{Log (Vascular LOX volume/}\mu\text{m}^3) = 9.5300615 - 1.1765328 * \text{Log (Tortuosity index)}$$
|  |  |  |  |  |
| --- | --- | --- | --- | --- |
| Mild/moderate CAA donors | 74 | -0.2287 | 0.0274 | ns |
| Severe CAA donors | 112 | ns |  |  |
| All CAA cases | 186 | ns |  |  |
| Neurological controls | 56 | ns |  |  |
| All cases | 242 | ns |  |  |

The enzyme LOX is a driver of vascular fibrosis and cross-links extracellular matrix polymer proteins to stiffen the connective tissue matrix of vessels ^31^. Its role in the brain is not well studied, although a few reports indicate it may be associated with β-amyloid pathology in Alzheimer’s disease and may be expressed in astrocytes ^32,33^. We hypothesized LOX activity may account for the increased vessel stiffness we observed in arterioles with CAA. A recent study using pan-lysyl oxidase inhibition in pancreatic cancer showed remodeling and crosslinking with fibrillar collagen matrices, reducing the tumor stiffness ^34^.

We produced a directly conjugated antibody for efficient staining of the tissue blocks and thoroughly validated LOX target specificity for immunostaining (supplementary fig. 2). We discovered that LOX was present in some β-amyloid plaques, as had previously been reported^35,36^. LOX was also present in vessels with CAA, primarily in the adventitia and especially in dilated, degenerating arterioles (fig. 4C). We observed that LOX was often not directly colocalized with β-amyloid but was nevertheless spatially related to β-amyloid deposits, and we aimed to quantify this relationship. We stained tissue blocks for collagen IV, methoxy-X04 (β-amyloid plaques), VSM, and LOX, analyzing controls, cases with mild/moderate CAA, and cases with severe CAA as in the previous section. Representative imaging results from the analysis across grades of vascular degeneration are shown in fig. 5. We quantified the relationship between the LOX staining and morphological features of degeneration, as shown in fig. 2 and 3. Vascular LOX was strongly related to β-amyloid abundance in the vasculature with Spearman correlation *(r_s_ (268) ρ= 0.5869 p<0.0001)*, fig. 6A-B. Vascular LOX was inversely related to the presence of VSM *(r_s_ (276) ρ= −0.4142 p<0.0001)*. LOX also correlated with vascular diameter *(r_s_ (231) ρ= 0.2854 p<0.0001)*, and the variability in vascular diameter *(r_s_ (231) ρ= 0.2759 p<0.0001)* (fig. 6D). Vascular tortuosity peaked with lower levels of LOX, but no statistically significant correlation was seen across the cohort. However, we found focal, asymmetric areas of LOX staining in arterioles were often present in twisted and distorted arteriole segments, and we hypothesized that asymmetric stiffening of the extracellular matrix may account for some of the arteriolar tortuosity observed in mild/moderate CAA, while heavier/circumferential areas of LOX reactivity have a symmetric effect on the vessel architecture (see fig. 6E and supplementary video 5).

**Figure 5:**
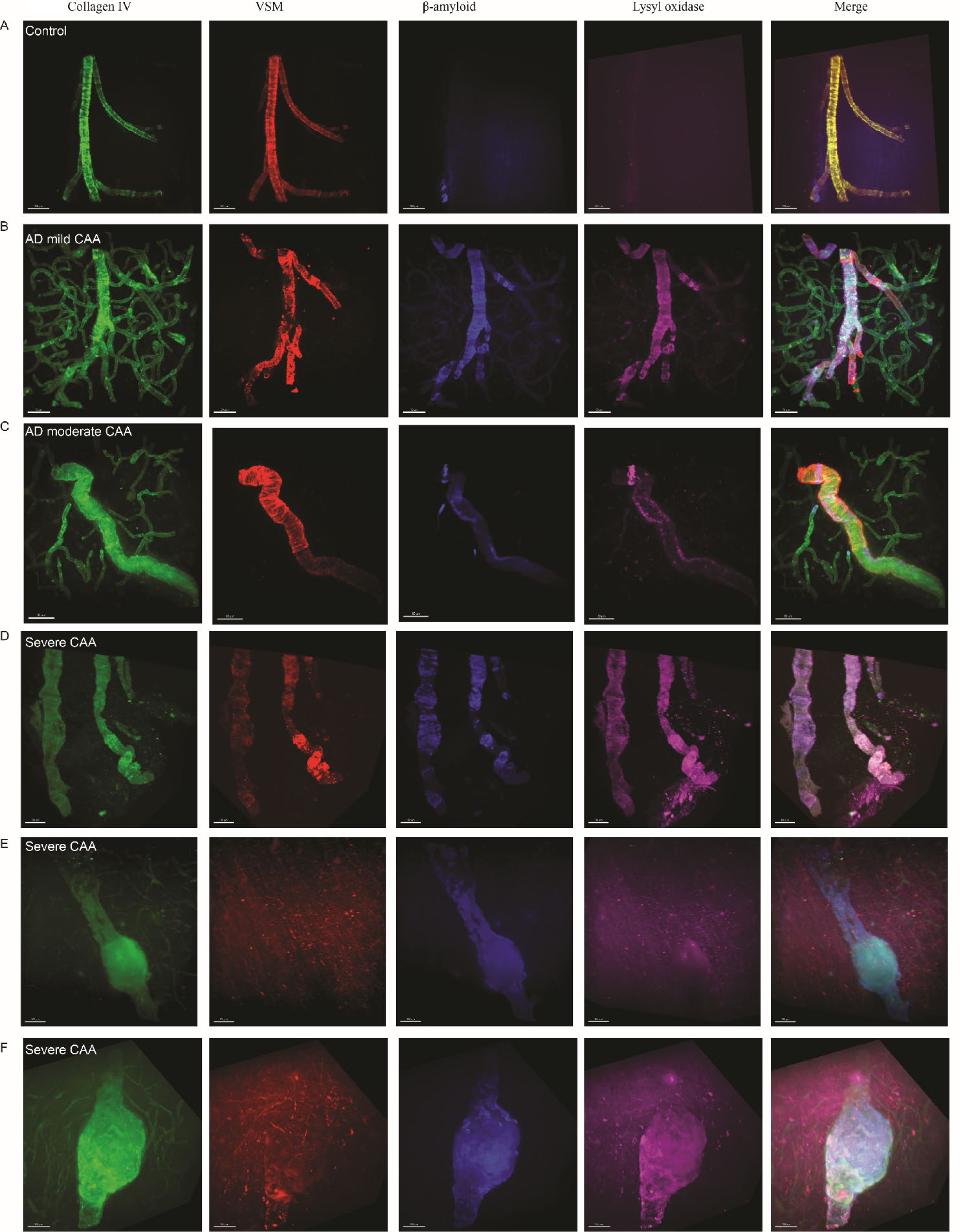
Morphological changes in vessels affected by CAA in relation to β-amyloid and LOX. Three-dimensional images of a control arteriole are compared with mild, moderate and severe CAA arterioles stained with Collagen IV-488 in green, VSM actin-Cy3 in red, methoxy-XO4 for β-amyloid, in blue and LOX-647 in far-red (shown in pink). A merged image of the 4 channels is shown in the last column, scale bar = 100μm. (**A**) Control with intact collagen IV and VSM, without methoxy-X04 or LOX. (**B)** Mild CAA shows a mild degeneration with patches of loss of VSM correlating with foci of β-amyloid and LOX staining. **(C)** AD with moderate CAA shows areas of degeneration of VSM with deposition of β-amyloid and LOX correlating with twisting distortions of the arteriole. **(D)** CAA with severe degeneration shown near-complete loss of VSM with heavy deposition of β-amyloid and LOX, tortuosity and dilation of the arteriole. **(E and F)** In severely degenerated arterioles with CAA the deposits of β-amyloid lose their organized ring shape and look fractured at sites with aneurysmal dilation of the arteriole.

**Figure 6:**
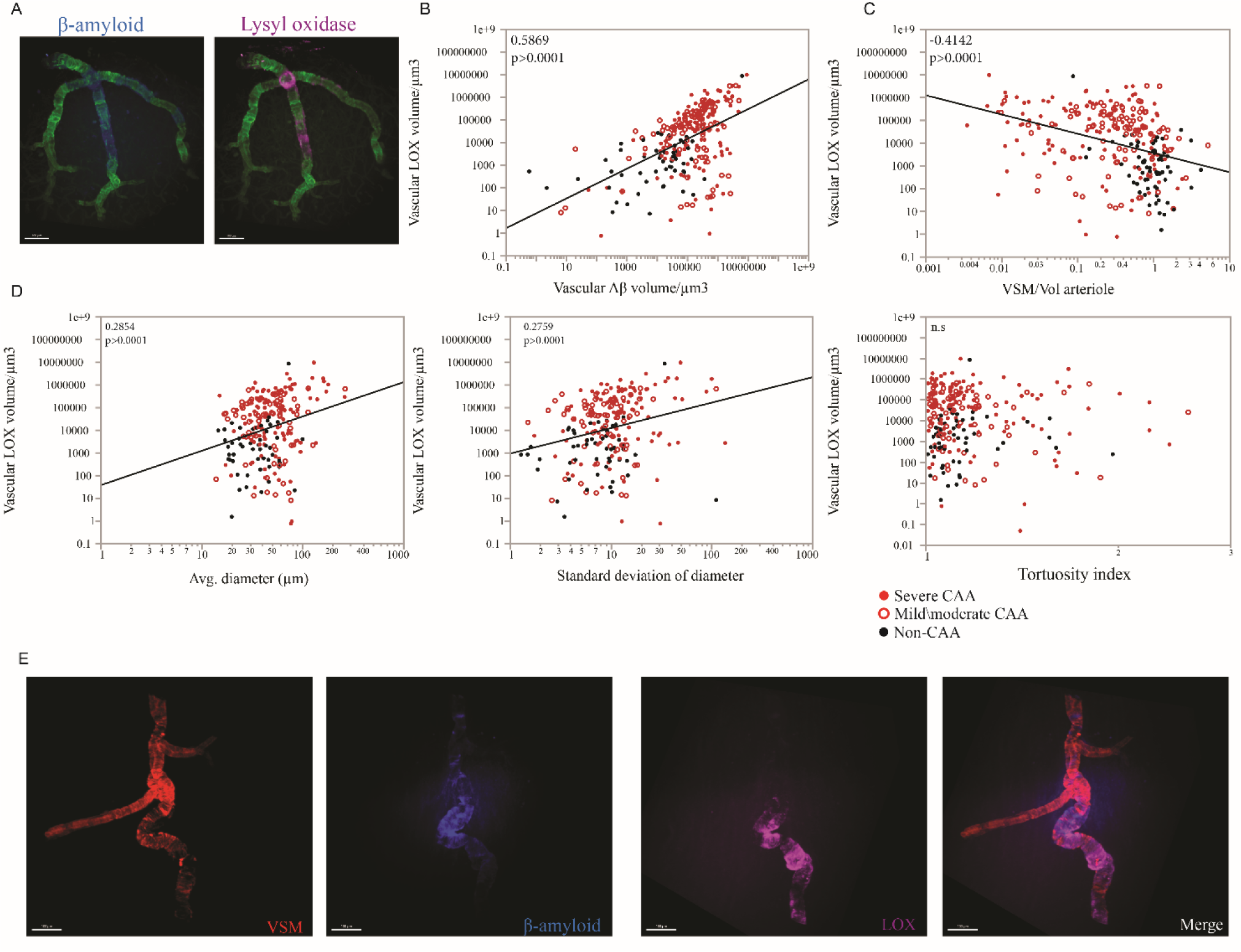
Morphological alteration of arteriolar walls correlates with volume of LOX. **(A)** β-amyloid in blue stained with Methoxy-X04 and LOX in pink, showing the overlap of β-amyloid and LOX on the arteriola, Scale bar with 100 µm. **(B)** The volume of LOX strongly correlated with vascular β-amyloid **(C)** and inversely correlated with vascular smooth muscle (VSM) volume. **(D)** LOX also correlated with increased vascular diameter and variability of the standard deviation of the diameter. While total volume of LOX did not correlate with tortuosity. **(E)** Eccentric deposits of LOX were frequently observed at sites of severe tortuosity. β-amyloid in blue stained with Methoxy-X04; VSM actin in red; LOX in pink, scale bar with 100 µm. See also supplementary video 5.

## Discussion

In this study we quantified morphological features of vascular degeneration in cerebral amyloid angiopathy. Degenerating arterioles were depleted of VSM and developed a widened lumen, likely due to laxity from the loss of VSM. We discovered that vessels with CAA were markedly stiffer than healthy vessels and this may be partly due to the effects of LOX in the extracellular space. At early stages of degeneration vessels were often tortuous which we attributed to the presence of focal, eccentric deposits of β-amyloid and/or LOX, while at later stages when both β-amyloid and LOX deposits tended to be extensive and concentric. Degenerating arterioles were also marked by dolichoectasia, areas of strictures and aneurysmal dilations. We also discovered that there is a close association between vascular β-amyloid deposition and cerebral microvascular degeneration including vessel wall rupture. Vessels associated with microhemorrhages were focally dilated near the site of hemorrhage with depletion of VSM and fractured shards of β-amyloid.

A major strength of this study is the use of quantitative three-dimensional microscopy. Neuropathology has traditionally relied on the study of thinly cut slices of brain tissue to enable adequate light penetration through the tissue for microscopy. Advances in confocal and multiphoton imaging have enabled better light penetration into tissue and consequently the ability to study thicker sections, but still significant tissue processing remains which can introduce artifacts and complicate the interpretation of microscopic findings. Three-dimensional tissue structure has historically been inferred by a variety of approaches, including the analysis of sequential tissue sections with digital reconstruction, but this approach is cumbersome and significant artifacts from tissue processing usually remain. This limitation is particularly problematic when trying to study networks (neuronal pathways or vascular networks) on a microscopic scale. The introduction of optical clearing techniques over the last decade or so, including most famously the CLARITY technique ^26^, has raised the possibility of studying relatively larger blocks of tissue rendered transparent for superior light penetration. Optical clearance of tissue for three-dimensional microscopy is regularly performed upon mouse brain tissue and other experimental model systems ^37-39^, but is rarely performed upon aged human brain tissue, and the few successful reports have mostly been on small (∼1mm thick) sections. We developed strategies to manage the high level of autofluorescence from accumulated pigments in brain tissue to leverage this innovative microscopy technique to study the cerebral microvascular network *in situ* on large (>1 cm^3^) tissue blocks from aged human brain.

In comparison to neuronal tau tangles and parenchymal plaques, the contribution of vascular pathologies to AD has been relatively understudied. Most patients with AD have some degree of CAA in autopsy studies, and 40% have some form of vascular cognitive impairment and dementia (VCID) ^40-43^. Behind AD, VCID is the second most common cause of dementia. CMH in a lobar distribution are a well-established clinical biomarker for CAA. CMHs are linked to β-amyloid burden on PET-amyloid imaging and to CSF markers of β-amyloidosis, and they predict both hemorrhage risk and deterioration of cognitive function over time ^44^. For this reason, the association of CMH with CAA is not in doubt. However, the mechanism of hemorrhage in CAA is incompletely understood. The most prevalent assumption is that β-amyloid is directly toxic to various elements of the arteriole leading to fragility and hemorrhage, and there is significant *in vitro* and *in vivo* evidence to support this view ^45^. However, previous reports that β-amyloid was not present at vascular rupture sites could question this hypothesis. An alternative hypothesis could be proposed that loss of VSM cells due to β-amyloid deposition in more proximal arterioles could lead to hyperperfusion from loss of autoregulation and turbulent flow due to disruption of the lumenal architecture, leading to vascular remodeling including distention and hemorrhage. Finally, it is possible that β-amyloid is directly toxic to the vessel at the site of hemorrhage, but activation of scavenging cells (before or after hemorrhage) clears the β-amyloid. Any or all of these mechanisms may be involved in the etiology of spontaneous hemorrhage in CAA, and identifying the major mechanisms will be critical to designing a treatment to prevent hemorrhages.

The studies questioning the association between vascular β-amyloidosis and cerebral microhemorrhage have problems that illustrate the inherent limitations of evaluating the microvasculature using two-dimensional techniques ^46,47^. In the cohort study by van Veluw and colleagues ^48^ three recent and four old cerebral microhemorrhages were found wherein the culprit vessel could be identified and β-amyloid was present in only one. In some cases, the authors interpreted hemorrhage products in the perivascular space as a CMH. We previously reported that hemorrhage from a CMH could propagate in the perivascular space some distance from the CMH, so in some of the cases it is likely that the tissue evaluated did not include the vessel-segment that ruptured ^49^. The study by Kovari and colleagues examined CMH in the white matter ^50^. It is well established that β-amyloid deposition in CAA is largely restricted to the cortical ribbon, so it is not surprising that the host vessel did not demonstrate β-amyloid deposition ^51^. Moreover, they interpreted microscopic perivascular hemosiderin deposits as CMH, but these are too small to produce the CMH detected by MRI and probably represent a distinct phenomenon. An earlier study by Fisher and colleagues found no deposition of β-amyloid around microhemorrhages in the putamen but CAA is not thought to be a typical cause of microhemorrhages in deep nuclei ^52^ which are usually from the sequelae of severe hypertension. Similar to the study by Kovari and colleagues, this study interpreted microscopic iron deposition around arterioles and capillaries as CMH. Unfortunately, the term CMH as it is used clinically is a misnomer because the hemorrhages are not in fact microscopic. We previously performed a series of studies to evaluate the size and iron content of “microhemorrhages” and to overcome the “blooming effect” that magnifies the apparent size of CMH on susceptibility weighted and gradient echo-T2* images. We found that iron concentration in CMBs was about 1.8 mg/g tissue (roughly 35 times the background iron level in neocortex), containing between 0.5 to 5 micrograms of iron in a typical microhemorrhage. This is equivalent to a detectable diameter of about 0.8 mm ^53-55^. Consequently, microscopic perivascular hemosiderin deposits are unlikely to be the pathological correlate of CMHs on MRI and the absence of β-amyloid deposition in vessels adjacent to these findings is not surprising. Nearly all CMH should be visible to the naked eye, with the possible exception of old, healed hemorrhages where hemosiderin laden macrophages are abundant and should be easily identified on microscopic examination. In the current study, we included subjects with severe CAA and numerous CMH and examined the tissue with three-dimensional imaging of optically cleared tissue which enabled confident identification of CMH and the culprit vessel. Our results support the hypothesis that β-amyloid may be directly involved in undermining the structural integrity of the vessel; although the other potential mechanisms may also play a role. Morphologically intact vessels with CAA had organized rings of β-amyloid which appeared to form around vascular smooth muscle fibers. When vessels become significantly dilated or rupture, these rings appeared to shatter, leaving behind sharply-contoured fragments or shards of β-amyloid. Animal studies have shown that heavy β-amyloid deposits cause cerebral arterioles to lose their pulsatility ^14^. We hypothesize that when these β-amyloid rings shatter in vessels depleted of VSM, the vessel may re-gain its pulsatility and the resulting movement of the sharply contoured shards within the vessel wall may tatter the connective tissue matrix leading to the latter stages of vascular degeneration. Because of the expansion (in some cases massive) in the arteriolar volume at rupture sites, the amount of β-amyloid in proportion to the vessel size may be fairly small, but it is consistently present.

A notable protein responsible for matrix remodeling and stiffening in blood vessels is LOX ^56^. LOX is necessary for blood vessel development as shown in morphogenesis studies in a mouse model ^57,58^. Atherosclerotic lesions contain pathological levels of LOX, and loss of LOX is associated with intracranial aneurysm rupture ^59-61^. Administration of irreversible pan-LOX inhibitor BAPN significantly diminished stiffness and tensile strength of aortic tissue in rats ^62^ We discovered that the volume of LOX on cerebral vessels strongly correlated with degenerative morphologies and β-amyloid deposition. Crosslinking of the extracellular matrix mediated by pathologic levels of LOX likely accounts in part for the increased vascular stiffness we observed in CAA. LOX is also a well-known driver of vascular fibrosis which could also contribute to increased vascular stiffness. A transgenic AD mouse model that overexpresses TGF-β1, another mediator of fibrosis and a possible regulator of LOX activity, has been shown to develop cognitive and vascular pathologies, lending support to the notion that this pathway is likely to be functionally important ^63^. There is an inherent limitation in neuropathological studies that it is difficult to establish causal relationships between the factors being studied. Consequently, our interpretation of the role of β-amyloid and LOX as playing key roles in vascular degeneration are informed by existing findings in the literature and will need to be confirmed in mechanistic studies. LOX catalyzes the oxidation of susceptible lysine residues, thereby producing highly reactive aldehyde groups that spontaneously form covalent bonds with oxidized and unoxidized lysine residues nearby leading to cross-linking of LOX substrates, such as collagen and elastin fibrils, which ensures structural stability of extracellular matrix ^64^. LOX activity is diminished as a result of impaired copper translocation in Menkes disease ^65^ because the insertion of copper in the nascent LOX is critical for its normal function. LOX is synthesized as a 48 kDa precursor of the proenzyme, or pre-pro-LOX ^66^. In the ER, the pre-pro-LOX is subjected to signal peptide cleavage at the N-terminus generating pro-LOX which is then N-glycosylated, yielding 50 kDa version that is secreted outside the cell. In the extracellular space, pro-LOX is cleaved by Bmp1 ^67,68^, yielding the mature 32kDa LOX ^69-71^. LOX is normally expressed in various human tissues; a northern blot detected higher RNA levels in kidney, placenta, pancreas, heart, and skeletal muscle and low levels in liver and the brain ^72^. In the brain, LOX is expressed primarily by astrocytes, VSM and endothelial cells ^68,73,74^. In addition to its role in vascular biology, LOX is also implicated in cell adhesion ^75^ and migration mechanisms in human disease, notably in hypoxia-induced tumor metastasis ^76,77^. LOX activity was reported to be increased in *post-mortem* brains of patients with AD and dementia compared to control subjects ^78^. Furthermore, LOX protein levels were found to be increased across 306 immunohistochemical studies of post-mortem AD brains, analyzed in a systematic review ^79^. In AD, LOX was spatially associated with β-amyloid plaques and vessels with β-amyloid in a rare hereditary form of CAA ^80^. In this study we extended these earlier observations by demonstrating a close association between the amount of LOX in vessels with CAA and microvascular degeneration. LOX is likely an important factor in vascular degeneration in CAA and these results strongly support further investigation of LOX and the LOX-family of enzymes as potential therapeutic targets for microvascular degeneration.

## Supplementary material

Supplementary methods figure 1:HeLa cells (American Type Culture Collection, United States) were grown in Dulbecco’s modified Eagle’s medium supplemented with 2 mM l-glutamine, 100 units/ml penicillin/streptomycin, and 10% (v/v) fetal bovine serum (FBS) at 37°C in an atmosphere of 5% CO2. Cells were plated in 6-well plates onto coverslip at a concentration of 1×106 cells/well and cultured overnight prior to transfection. Cells were transiently transfected with 1 µg of pCMV-human LOX-GFPS (HG17796-ACG; Sino Biological) using FuGENE® Transfection Reagent according to the manufacturer’s instructions. For the immunocytochemistry cells with/without transfection were fixed in 4% paraformaldehyde, washed with TBS containing 20 mM glycine, permeabilized with 0.1% TritonX-100 and labelling LOX with NB100-2527 (2 µg/mL) at 4°C overnight. After cells were washed and stained with Donkey Anti-Rabbit IgG H&L Alexa Fluor®488 or Alexa Fluor®594 at 1:1000 for 1h at room temperature. Nuclei were stained with DAPI and after extensively washed cells were mounted in a Vectashield immunofluorescence medium (Vector Laboratories, United States). Microscope observations were performed with a 63x oil immersion objective using a Zeiss LSM 710 Confocal Microscope (Zeiss, United States).

### Supplementary video

Video 1: This video showcases the methodology for image reconstruction of immunological staining with vascular markers on a representative vessel with CAA pathology. Collagen IV in green is used as a vascular marker. Vascular smooth muscle actin (VSM) is stained in red. Notably, VSM is missing in the areas of β-amyloid deposition, stained with methoxy-X04 in blue. The reconstruction shows a strong correlation between the loss of VSM volume and β-amyloid deposits. Scale bar 50 µm.

Video 2: The video contrasts the vascular networks of a subject with normal cerebrovascular network and two patients with cerebral amyloid angiopathy. The prominent morphology between the cases include a number of dilated arterioles, many with irregular lumenal diameter and some with increased tortuosity compared to the healthy case. Several microhemorrhages are also visible in the images. Three-dimensional imaging was performed on a SmartSPIM single-plane illumination light sheet microscope (LifeCanvas Technologies) with a 3.6× objective (NA = 0.2) and axially swept rolling shutter. Scale bar 200 µm.

Video 3: The video displays a vessel with CAA and contrasts areas of β-amyloid deposits which have organized ring-shaped morphology with an area of dilation in the vessel wall where the β-amyloid rings appear to have fractured with sharply contoured “shard-like” morphology. β-amyloid is stained blue with Methoxy-X04, while the blood vessels are stained with collagen IV-488. Scale bar 50 µm.

Video 4: The video provides examples of ruptured vessels associated with microhemorrhages. Shards of β-amyloid are present at or near the sites of rupture. Three-dimensional images of arterioles are shown with staining of β-amyloid using thiazine red (red) and lectin in green to label the vessel wall. The last two images show vascular surface rendering from ruptured vessels. Scale bar of varied length at the bottom left.

Video 5: The video illustrates the relationship between LOX and vascular degeneration in CAA. Lysyl oxidase is stained in pink, collagen IV in green, smooth muscle actin (SMA) labeling vascular smooth muscle in red, and β-amyloid in blue. The reconstruction highlights the substantial spatial positive correlation of β-amyloid deposition with lysyl oxidase and the negative correlation with VSM. Scale bar of varied length at the bottom left.

---

**Supplementary figure 1:**
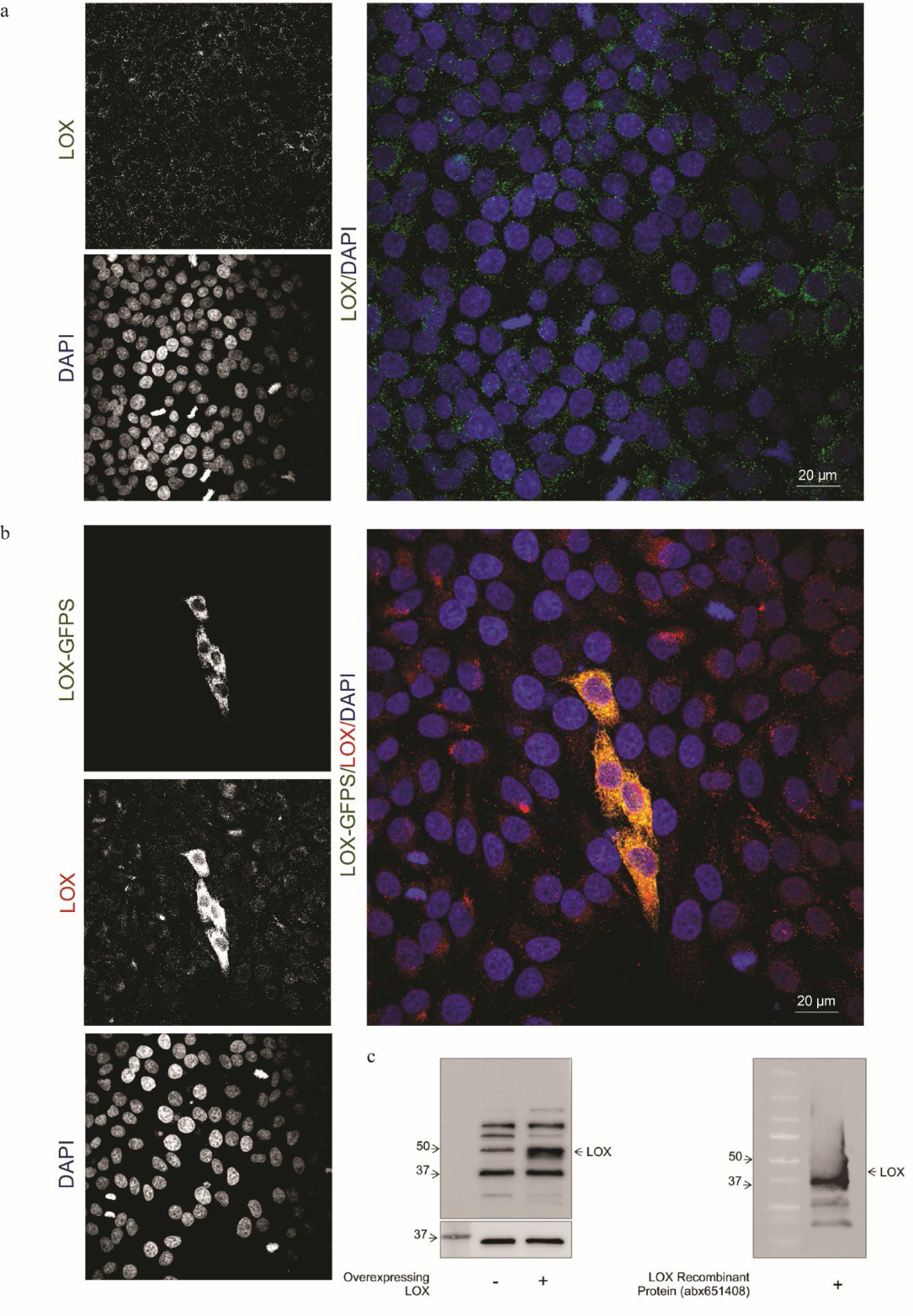
Immunocytochemistry validation of LOX antibody (Novus; NB100-2527) in HeLa cells. Cells were fixed, permeabilized and stained for LOX followed by Donkey Anti-Rabbit IgG H&L Alexa Fluor®488 or Alexa Fluor®594. (a) Image shows endogenous LOX levels (green). (b) Counterstained image shows immunodetection of LOX (red) in cells that were transfected with pCMV-LOX-GFPS (green). Yellow/orange fluorescence indicate areas where the antibody recognized the fluorescence labelled LOX. Images are of 5 μm z-stack scanning with a step interval of 1 μm, n=3. Nuclei were stained with DAPI. The scale bars are indicated. (c) Hela cells were transfected with 2 ug of LOX plasmid pCMV3-LOX (HG17796-UT, Sino Biological) using the guidelines FuGENEâ HD protocol, then the cell proteins were extracted and 20 ug of cell lysates were denatured, separated, and transferred via Western Blot following the protocol previously described. To check the LOX protein expression was used Rabbit anti-LOX antibody (1 ug/ml) (NB100-2527, NOVUS) and like a loading control Mouse anti-GAPDH antibody (1 ug/ml) (ab59164, Abcam). The images were obtained using secondary fluorescence antibodies (1:2500) (LICOR) in the Chemidoc MP Imaging System (BIO-RAD). Next, image with 5 ug of LOX recombinant protein (abx651408, abbexa) were denatured, separated, and transferred using Western Blot protocol previously described. To check the LOX protein expression was used Rabbit anti-LOX antibody (0.5 ug/ml) (NB100-2527, NOVUS). The images were obtained using Goat anti-Rabbit-HRP secondary antibody (100 ng/ml) (65-6120, Invitrogen) and SuperSignalä West Femto Maximun Sensivity Substrate in the Chemidoc MP Imaging System (BIO-RAD).

